# Histopathological aspects of induced resistance by *Pseudomonas protegens* CHA0 and β-aminobutyric acid in wheat against *Puccinia triticina*

**DOI:** 10.1101/2020.02.06.934943

**Authors:** Fares Bellameche, Mohammed A. Jasim, Brigitte Mauch-Mani, Fabio Mascher

## Abstract

After perception of specific biotic or abiotic stimuli, such as root colonization by rhizobacteria or selected chemicals, plants are able to enhance their basal resistance against pathogens. Due to its sustainability, such induced resistance is highly valuable for disease management in agriculture. Here we study an example of resistance against wheat-leaf rust induced by *Pseudomonas protegens* CHA0 (CHA0) and β-aminobutyric acid (BABA), respectively. Seed dressing with CHA0 reduced the number of sporulating pustules on the leaves and the expression of resistance was visible as necrotic or chlorotic flecks. Moreover, a beneficial effect of CHA0 on growth was observed in wheat seedlings challenged or not with leaf rust. BABA was tested at 10, 15 and 20 mM and a dose-dependent reduction of leaf rust infection was observed with the highest level of protection at 20 mM. However, BABA treatment repressed plant growth at 20 mM. Balancing between BABA-impact on plant growth and its protective capacity, we selected 15 mM as suitable concentration to protect wheat seedlings against leaf rust with the least impact on vegetative growth. To understand the mechanisms behind the observed resistance, we have studied the histological aspects of the fungal infection process. Our results showed that the p re-entry process was not affected by the two resistance inducers. However, both treatments reduced fungal penetration and haustoria formation. The timing and the amplitude of the resistance reactions was different after bacterial or chemical induction, leading to different levels of resistance to leaf rust. During fungal colonization of the tissues, a high deposition of callose and the accumulation of H_2_O_2_ in both CHA0-and BABA-treated plants pointed to an important contribution to resistance.

## INTRODUCTION

Plants dispose of several layers of sophisticated defense mechanisms to defend themselves against pathogen attack. The first layer is given by preformed physical and chemical barriers that impede the pathogen to penetrate into the plant and to initiate infection (Ferreira *et al.* 2006). Once the presence of the pathogen has been detected, the plant activates further chemical and physical barriers that block or at least delay the attack (second layer; Jones and Dangl (2006)). Defense success depends on the readiness of the plant to detect the pathogen. In the case of the interaction between wheat and the leaf rust pathogen (*Puccinia triticina*), the plant can detect specific fungal avirulence factors (elicitors) with leaf rust resistance genes (Lr). This gene-by-gene interaction is a very rapid recognition-reaction event leading to an elevated degree of resistance against the disease. However, the avirulence patterns can change and the pathogen may become undetectable by the plant. This resistance breakdown happened recently with yellow rust (Hovmøller *et al.* 2010) and stem rust (Singh *et al.* 2011).

In the case of unspecific recognition of the pathogen, the plant is still able to contain the development of the pathogen but with a reduced and variable degree of severity of infection (Jones and Dangl 2006). The degree of this quantitative resistance is linked to the readiness of the plant defenses and depends on a series of genetic and environmental factors. Besides the pathogen itself, biological and abiotic stimuli as well as certain chemicals can enhance plant resistance (Mauch-Mani *et al.* 2017). Such induced resistance can be limited to the site of the inducing treatment but it can also be systemic and thereby effective in parts of the plant distant from the site of induction (Van Loon 1997). For instance, certain root-associated bacteria such as the biocontrol strain *Pseudomonas protegens* CHA0 (formerly *P. fluorescens* CHA0) induce systemic resistance against viral and fungal diseases in various dicots (Maurhofer *et al.* 1994; Haas and Keel 2003; Iavicoli *et al.* 2003) and monocots (Sari *et al.* 2008; Henkes *et al.* 2011). Certain chemical compounds can also induce disease resistance in plants, *e.g.* the non-protein amino-acid β-amino-n-butyric acid (BABA). Root colonizing bacteria and BABA root treatment reduce significantly the severity of infection caused by the oomycete *Hyaloperonospora arabidopsis* on *Arabidopsis thaliana*, and the induced state is regulated by different defense signaling pathways, depending on the inducing agent and the challenging pathogen (Van der Ent *et al.* 2009).

In the present work, we aimed to study the mechanisms underlying induced resistance by CHA0 and BABA in wheat against leaf rust. A previous study has shown that root colonization by *Pseudomonas protegens* strain CHA0 reduces the number of leaf rust uredia on susceptible wheat seedlings in wheat (Sharifi-Tehrani *et al.* 2009). The enhanced resistance is very likely due to a resistance priming event by <u>i</u>nduction of <u>s</u>ystemic <u>r</u>esistance (ISR). This priming enables the plant to cope with the pathogen at an early stage of infection. To study this, in the work presented here, we followed the interaction between the plant and the pathogen at the microscopic level (De Vleesschauwer *et al.* 2008).

The infection process of leaf rust is well known (Bolton *et al.* 2008). After adhesion of a urediniospore on the leaf surface, germination, directed growth of the germ tube on the plant surface towards a stoma, and recognition of the guard cell lips take place. A small appressorium is formed over the stomatal opening and then, a penetration hypha is entering through the stomatal pore. Following penetration, a substomatal vesicle, and haustorium develop (Bolton *et al.* 2008).

Primed plants recognize the pathogen and produce reactive oxygen species (ROS) and deposit callose at the infection sites (Balmer *et al.* 2015). This rapid local oxydative burst generates, among other, hydrogen peroxide (H_2_O_2_) during pre-haustorial resistance against wheat leaf rust caused by *P. triticina* (Wesp-Guterres *et al.* 2013; Serfling *et al.* 2016). Callose is an effective barrier that is induced at the sites of attack during the early stages of pathogen invasion (Luna *et al.* 2011). A strong deposition of callose has been reported for the wheat Thatcher near-isogenic lines carrying leaf rust resistance genes (Wang *et al.* 2013).

Few studies have investigated rhizobacteria-and BABA-induced resistance against wheat leaf rust. In this study, we aimed to compare mechanisms involved CHA0-ISR and BABA-IR during interaction between leaf rust and wheat. To this end, we evaluated microscopically the development of fungal structures, the occurrence of callose deposition and hydrogen peroxide accumulation in leaf tissues.

## MATERIALS AND METHODS

### Induced resistance assay

#### Plant material and growth conditions

Experiments were done with the leaf rust-susceptible bread wheat cultivar Arina (Agroscope/DSP). Surface sterilized seeds were used in all experiments. Wheat seeds were rinsed with 70% ethanol, incubated for 5 minutes in 5 % bleach (sodium hypochlorite solution, Fisher Chemical, U.K.) and washed three times in sterile distilled water. The sterilized seeds were germinated on humid filter paper (Filterkrepp Papier braun, E. Weber & Cie AG, 8157 Dielsdorf, Switzerland) in plastic bags maintained in the dark at room temperature. Three to 4 days later, seedlings at similar growth state and morphology were selected and planted in 120 mL polypropylene tubes (Semadeni, 3072 Ostermundingen, Switzerland) filled with a standard potting mixture (peat/sand, 3:1, vol/vol). The tubes were placed in a growth chamber with the 16 hours day at 22°C and 8 hours night at 18°C and with 300 μmol m^−2^ s^−1^ light. The plants were watered regularly keeping the potting soil wet yet avoiding its saturation.

#### Bacterial inoculum

The bacterial inoculum consisted of the biocontrol agent *P. protegens* strain CHA0-Rif (Natsch *et al.* 1994) (in the following called CHA0), a spontaneous rifampicin resistant strain of *P. protegens* strain CHA0 (Stutz *et al.* 1986; Ramette *et al.* 2011). Both strains are similar in terms of growth rates, production of antimicrobial compounds (Natsch *et al.* 1994) and their capacity to induce resistance in wheat (Sharifi-Tehrani *et al.* 2009). Routinely, the strain was grown on solid King’s medium B (Pseudomonas agar F, Merck KGaA, 64271 Darmstadt, Germany) supplemented with 50 μg mL^−1^ rifampicin at 25°C in the dark for 3 days. For long-term storage, 1mL of a freshly grown bacterial suspension in King’s liquid medium B (30g proteose-peptone, 1.5g K_2_HPO_4_, 2.46 g MgSO_4_, 1.5g glycerol in 1 L distilled water) was mixed with 1mL glycerol (87%) and conserved at −80°C. For inoculum production, a single colony of a freshly grown culture was transferred to a 300 mL Erlenmeyer flask filled with 100 mL of King’s liquid medium B supplemented with 50 μg mL^−1^ rifampicin. After 12 h incubation at 28°C with continuous shaking at 150 rpm, the bacterial culture was centrifuged at 3700 rpm and washed twice with sterile 10mM MgSO_4_ solution. The final pellet was re-suspended in 20 mL sterile distilled water and adjusted to an OD_600_ of 0.1 corresponding to approximately 10^6^ CFU/mL and used for seed inoculation. For this, the sterilized wheat seeds were immersed in the bacterial suspension for 6 hours with shaking at 35-40 rpm at room temperature. Inoculated seeds underwent the pre-germination procedure as described above. Control seeds were soaked in distilled water for the same time period before pre-germination.

#### Treatment with β-aminobutyric acid

The resistance inducer BABA was purchased at Sigma-Aldrich (Buchs SG, Switzerland). Dilutions of 10, 15 and 20 mM of BABA in distilled water were used as a soil drench. For this, 10 ml of BABA solution were added to the soil to plants were at the 2 leaf stage, 48 hours before infection with leaf rust. Control plants were treated with the same amount of distilled water.

#### Effect of CHA0 and BABA on plant development

In a first step, the impact of the inoculation of CHA0 and BABA treatment on the plant was assessed. To measure root colonization by CHA0, 0.1g each of inoculated or control roots were shaken each in 10 mL sterilized distilled water during 1 min on a benchtop vortex mixer, followed by 1 min of sonication. The root extract was serially diluted and plated on solid King’s medium B supplemented with 100 μg mL^−1^ of rifampicin. The plates were incubated at 28°C in the dark and the number of CFUs was determined after 24h to 36h.

To investigate possible effects of CHA0 and BABA treatments on plant growth, the dry mass of the shoot of pre-treated seedlings was measured at 12 days after inoculation with leaf rust. Shoot length was defined as the upper part of the plant cut at the residue of the seed. The shoots were weighed (fresh weight), placed on coffee filter paper and dried separately in an oven at 65° until sample weight remained constant (dry weight).

#### Inoculation with P. triticina

Inoculations with leaf rust (*P. triticina*) were done at the 2-leaf stage (BBCH 12 (Meier 1997)) using freshly harvested urediniospores of isolate Pr2271 (Agroscope, Changins, Switzerland). The urediniospores were generated on leaves of cv. Arina. For infections, fresh urediniospores were mixed with talcum powder in a 1:9 w/w ratio and rubbed gently on the leaf surface. Inoculated plants were placed in a dew box in the dark at 18 to 22°C for 24h to promote infection. Subsequently, the plants were placed in the growth chamber as described above. After 12 days or when the symptoms were sufficiently developed on the control plants, the infection type was assessed using the 0–4 scoring system (Table S1) described by Roelfs (1992).

### Histochemical assessment of leaf rust infection in presence of CHA0 and BABA

#### Assessment of fungal growth and development

Leaf rust growth was observed on 2 cm leaf segments from the centre of the second leaves at 0, 6, 12, 24, 48, 72 and 96 hai (hours after inoculation). The leaf segments were immerged in 96% ethanol for 2-3 days to remove chlorophyll. The distained leaf segments were washed in an ethanol/ water (1:2 v/v) solution and then incubated in 0.5 M sodium hydroxide for 15 min with slight shaking. The leaf segments were incubated for 15 min in distilled water and before soaking for 2 h in 0.1 M Tris–HCl buffer (pH 8.5). Fungal structures were then stained with a 0.2% Calcofluor White solution in water (Sigma-Aldrich, Germany) for 5 min. After four washings in distilled water, the samples were stored in 50 % (v*/*v) glycerol for microscopic observation.

The preparations were examined with an epifluorescence microscope (Model E800; Nikon Instruments Europe, Badhoevedorp, The Netherlands) using excitation at 365 nm in combination with a 450 nm barrier filter and a dichroic mirror at 400 nm. This installation allowed the determination of position and number of all fungal organs on and in the leaf, namely germinated and non-germinated spores, appressoria, sub-stomatal vescicles and haustoria.

#### Identification and quantification of callose deposition

Assessment of callose deposition was done on segments from the centre of the second leaf at 0, 24, 48 and 72 hai with leaf rust according to Scalschi *et al.* (2015). The leaf tissue was distained for 48h in 96% ethanol until transparent. Subsequently, the leaf tissue was rehydrated in 0.07 M phosphate buffer (pH =9) for 30 min and incubated for 15 min 0.05% aniline-blue (Sigma, St. Louis) prepared in 0.07 M phosphate buffer and were finally stained overnight in 0.5% aniline-blue microscopic observations were performed with the epifluorescence microscope using a UV filter as described above.

The presence and the quantity of deposited callose was determined from digital photographs by counting the number of white pixels (representing callose deposits) in 20 infection sites for each replicate, using the GNU Image Manipulation Program (GIMP 2.10.10) software. Contrast settings of the photographs were adjusted to obtain an optimal separation of the callose signal from the background signal. Callose was automatically identified using the “Color Range” tool and callose-corresponding pixels were recorded as the area covered by the total number of selected pixels (Scalschi *et al.* 2015).

#### Accumulation of H_2_O_2_ at the infection sites

Detection of H_2_O_2_ was carried out using DAB (3,3-diaminobenzidine, Sigma-Aldrich, Switzerland) staining as described (Thordal - Christensen *et al.* 1997). The second fully expanded leaves were cut at 0, 24, 48 and 72 hai and immediately immersed in a solution containing 1mg mL^−1^ DAB dissolved in HCl acidified distilled water (pH 3.8). Leaves were incubated in the dark for 8 h to allow DAB uptake and reaction with H_2_O_2_. Subsequently, leaves were cleared in saturated chloral hydrate and scanned at 1.200 dpi (Epson perfection, V370 PHOTO).

In presence of H_2_O_2,_ DAB is reduced to a dark-brown deposit that can be easily visualized in the leaves. The H_2_O_2_ content of the leaves was quantified by counting the number of dark-brown DAB pixels using GIMP 2.10.10 software and the percentage of DAB stain was calculated corresponding to total leaf area (Luna *et al.* 2011). The dark-brown DAB pixels were selected using “Color selection” and the total area of leaves was using the “Free Selection” tool.

### Experimental set up and statistical analyses

All experiments were repeated at least twice. The induced resistance assay consisted of seven biological replicates. The fungal growth and the callose deposition assessments were done with three independent replicates and the H_2_O_2_ quantification on ten biological replicates. Fungal development structures were identified and counted at 50 sites in each replicate. The percentage of germinated spores = (germinated spores/observed spores)×100, the percentage of stomatal appressoria = (stomatal appressoria/germinated spores)×100, percentage of sub-stomatal vesicles = (sub-stomatal vesicles/stomatal appressoria) x 100 and percentage of haustoria = (haustoria/ sub-stomatal vesicles ×100) were determined.

Data were collected and stored in spreadsheets (Microsoft^®^ Excel 2010, Redmond USA). Statistical analysis was conducted in R (R Core Team, 2017).

In all experiments, statistically significant differences in response to CHA0 and BABA treatment compered to control were tested with Student’s T-test, Except, the experiment data of the effect of CHA0 and BABA on plant development were analyzed by two-way ANOVA with the factors; treatment (CHA0 and BABA) and rust inoculation (infected or not). Tukey Honest Significant Differences (HSD) test was used for multiple comparisons. Significant differences were considered at *P* <0.05.

## RESULTS

### Plant growth and biomass production in presence of CHA0, BABA and following inoculation with leaf rust

Twelve days after planting of the seedlings, 5×10^5^ CFU/g on average of CHA0 were recovered on the fresh roots, showing the capacity of the bacterium to successfully colonize the roots. In preliminary experiment, the initial concentration of bacteria (10^4^, 10^6^ and 10^8^ CFU/mL) used for seed inoculation did not alter the final number of bacteria on the roots (data not shown).

The effect of CHA0 and BABA on plant length and biomass production is presented in Fig. 1. The results indicate that the plants treated with CHA0 grew significantly longer and produced significantly more biomass. The growth and the biomass were not influenced by the presence of the pathogen (Fig. 1A and B). In contrast, plants treated with BABA at 20 mM were significantly shorter and produced significantly less biomass than the untreated control. Meanwhile, the plants treated with 10 or 15mM of BABA were not different to the untreated control. The infections with leaf rust did not affect plant growth or biomass, exception made, for the treatment with BABA at 20 mM (Fig. 1C and D).

**Figure 1:**
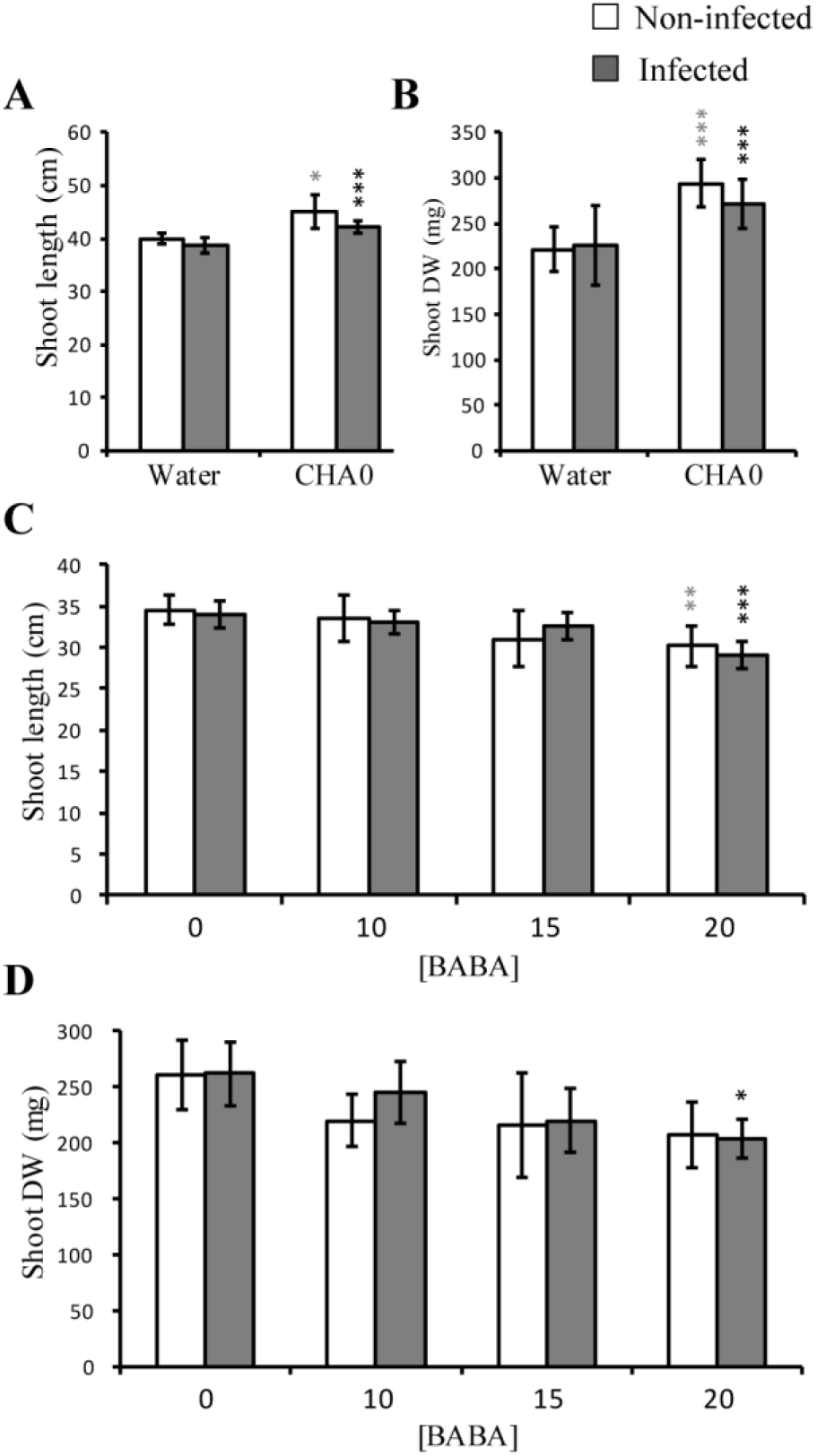
Growth and biomass production of plants treated with CHA0 (**A** and **B**) and BABA (10, 15 and 20 mM) (**C** and **D**) at 12 dpi with *P.triticina.* Shoot length was measured from the seed to the top of the longest leaf. Shoots dry weight was obtained after drying samples at 65° until the weight remained constant. Error bars indicate the standard errors for the average values of 7 replicates, grey and dark stars indicate significant differences compared to non-infected and infected control respectively (Tukey’s test; P < 0.05).

### Phenotypic reaction to leaf rust of seedling pre-treated with CHA0 or BABA

Twelve days after inoculation with *P. triticina*, control plants were totally healthy (Fig. 2a) while the rust inoculated plants presented uniformly uredia with chlorosis corresponding to score 3 (high infection type (HI) (Roelfs 1992)) (Fig. 2b). When treated with CHA0 (Fig. 2c), leaves showed overall a lower number of uredia compared to the infected control. The symptoms are heterogeneous, namely chlorotic flecks (score “;”, low infection type (LI)) but also uredia without sporulation (score “2”, LI) and with sporulation and a chlorotic halo (score “3”, HI).

**Figure 2:**
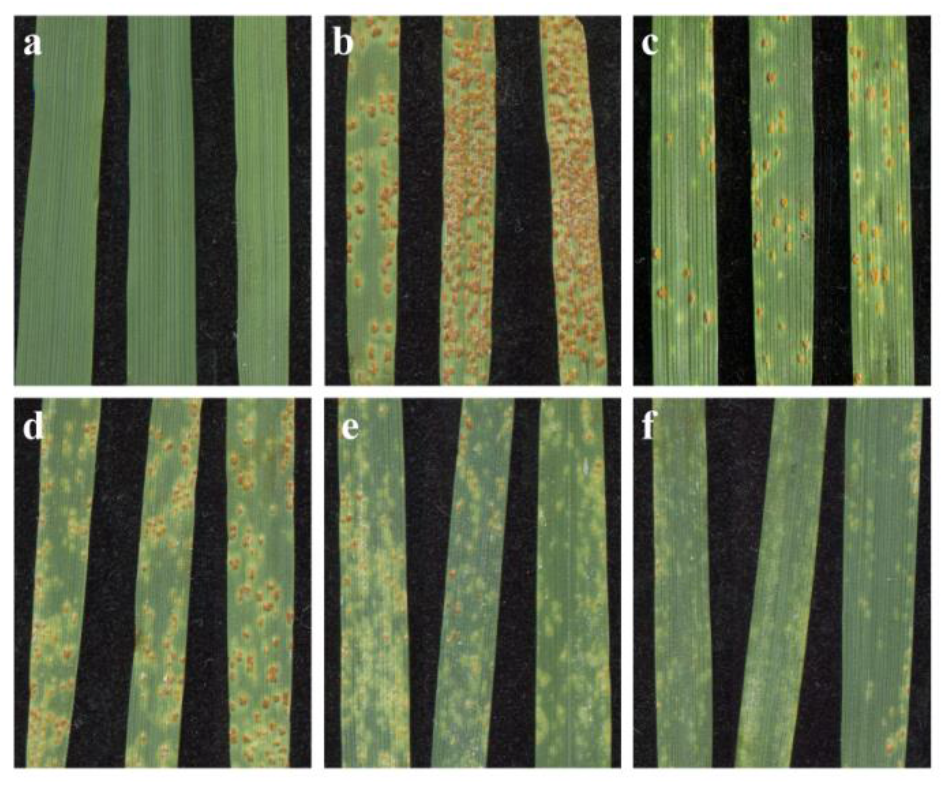
Leaf rust infection on seedling leaves of cultivar Arina at 12 dpi. **a,** control plant non-infected, **b,** infected plants non-treated, **c,** infected plants pre-treated with CHA0, from **d** to **f** infected plant treated with different concentration of BABA 10, 15 and 20 mM, respectively. Images were obtained by scanning at 1.200 dpi a segment of 3 to 4 cm from the centre part of the second leaf.

Similarly, BABA treatments also resulted in a mix of chlorotic flecks (score “;”) and small to medium pustules with and without low sporulation scored as “1” and “2”. Generally, all BABA treatments lead to low infection type symptoms. Yet, the scores were clearly dose-dependent, since the higher the BABA concentration, the lower the scores (Fig 2d, e, f).

### Fungal infection structures

Calcofluor white staining was used to track the pathogen structures during the first 96 hours after infection (hai) with leaf rust in non-treated control and on the plants pre-treated with CHA0 and BABA at 15mM. Within 6 hai, germ tubes started to elongate (Fig. 3A, 1). Independent of the pre-treatment, about 90% of urediniospores had germinated within 6 hai, in all treatments (Fig. 3B, 1). In the following, the proportion of germinated spores remained constant.

**Figure 3:**
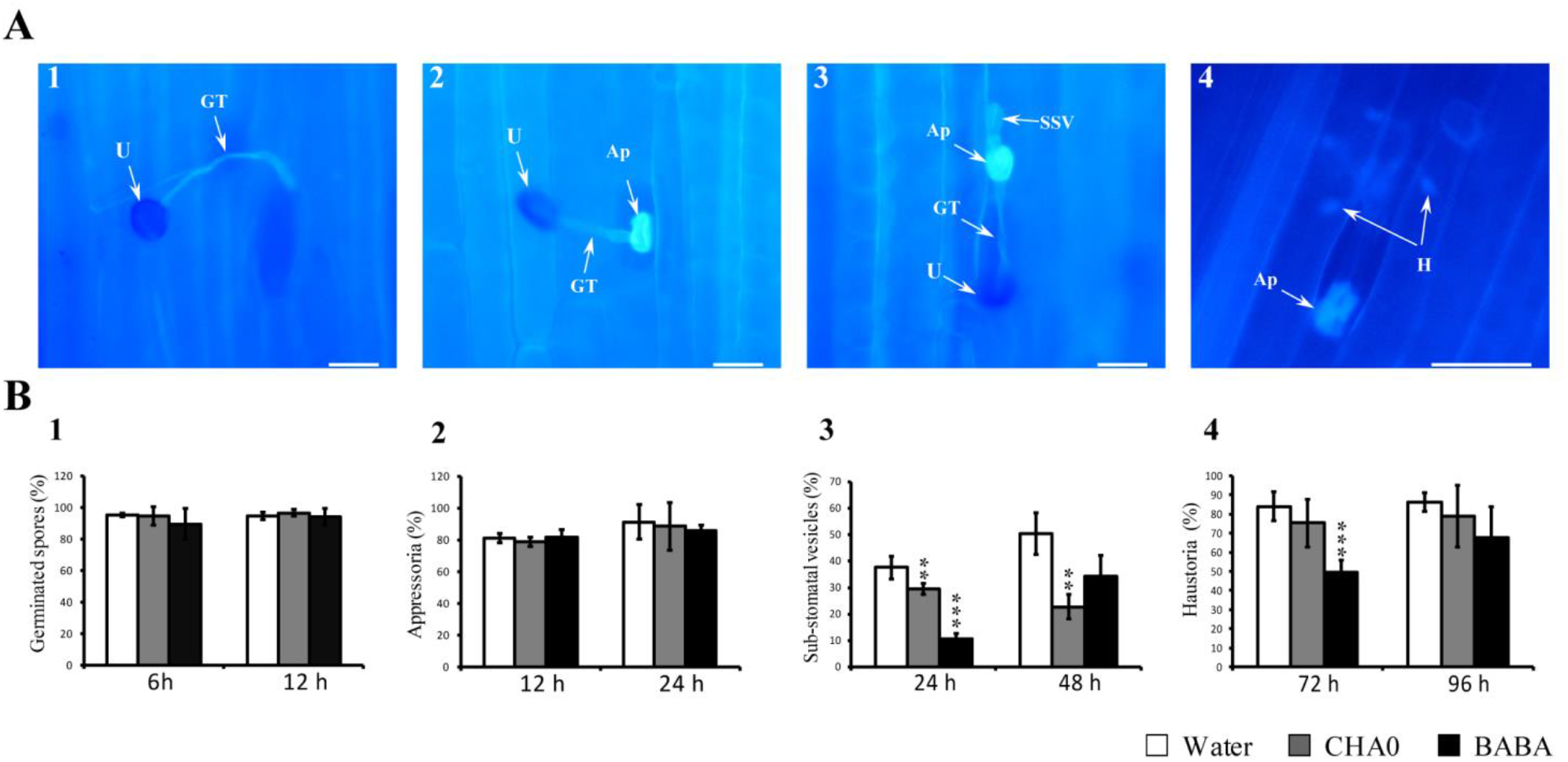
Microscopic observations and quantification of fungal structures *P. triticina* in wheat seedlings. **A**, fungal structures stained with calcofluor white and visualized under the epifluorescence microscope. In **B,** Percentages of fungal infection structures during infection of wheat with *P. triticina*: (1) spores germination, (2) appressoria, (3) sub-stomatal vesicles and (4) haustoria. Treatments: **CHA0**, plants obtained from seeds inoculated with CHA0 (10^6^ CFU/ml), **BABA**, plants soil-drenched with BABA (15 mM) 48h before rust infection, **Water**, plants mock-treated with sterile distilled water. Fungal structures: **U**, urediniospore, **GT**, germ tube, **Ap**, appressorium, **H**, Haustoria. Error bars indicate the standard error of the average values of 3 replicates at 50 infection sites for each replicate. Asterisks indicate statistically significant differences in response to CHA0 or BABA treatment (Student’s *t*-test; P =0.05). Bar 20 μm.

Once germinated, the fungus forms appressoria at the stomatal regions (Fig. 3A, 2). The formation of appressoria started at 6 hai (data not shown). Yet, at 24 hai, 85-88 % of the germinated spores had formed appressoria (Fig. 3B, 2). This proportion hardly varied between the control and the bacterial and BABA treatments.

Through the appressoria, the fungus penetrated into the cavities below the stomata, forming infectious vesicles in the substomatal cavity (Fig. 3A, 3). Our observations indicate that the formation of those vesicles started at 12 hai (data not shown). On the leaves of non-treated control plants, about 37% of appressoria had formed vesicles after 24 hai, with the proportion increasing to 50% after 48 hai. In plants inoculated with CHA0, about 29% of the appressoria formed. Yet, after 48 hai, the proportion of formed vesicles was about 23%. In BABA (15mM) treated plants, the proportion of formed vesicles was 10% and increased to 30% at 48 hai (Fig. 3B, 3). At 48h we noticed the formation of haustoria out of the vesicles (Fig. 3A, 4). At 72 hai, more than 80% of the sub-stomatal vesicles had formed haustoria in the untreated control plants. This proportion did not change at the last time point, at 96 hai. In the CHA0 treated plants, the proportion of formed haustoria was not different to the control plants, at both time points (i.e. 72 and 96 hai). However, the absolute number of haustoria was significantly lower in the CHA0 treated plants compared to the control. With BABA treatment, only about 50% of the vesicles formed haustoria. This is significantly less haustoria formed than in the control. At 96 hai, haustoria formation increased in the BABA treatment to 70% and there was no significant difference with the other treatments. Also, in the BABA treatment, the absolute number of haustoria was lower compared to the control plants (Fig. 3B, 4).

### Callose deposition after leaf rust inoculation

Callose was quantified at 24, 48 and 72 hai after inoculation with leaf rust in the control, the CHA0 and the BABA 15mM treatment (Fig. 4). Callose deposition at the infection sites was made visible by aniline bleu. The amount of deposited callose was measured by counting pixels of stain around the infection sites (suppl. Fig. 1S). Overall, callose deposition occurred in all treatments within the first 24 hai (Fig.4). However, in plants pre-treated with CHA0 and BABA, a significantly higher quantity of callose was observed compared to the control. With CHA0, callose accumulated at the guard cells and was highest at 72h hai. In plants treated with BABA, we measured the highest callose deposition after 48 hai. Here, callose was not only observed at the guard cells (stomata) but also in mesophyll cells, at 72h.

**Figure 4:**
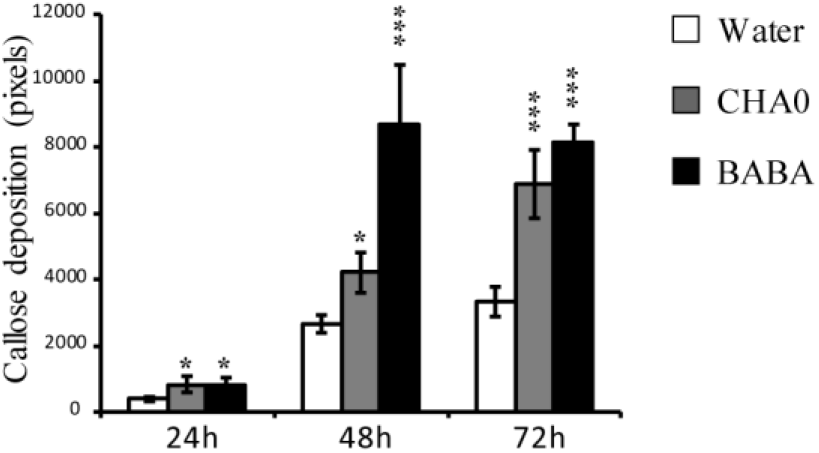
Callose deposition in wheat leaves in response to *P. triticina* infection in treated and control plants at 24, 48 and 72 hai. Callose was identified as white spots around the infection sites and quantified from digital photographs by measuring the number of pixels of the white surface at 20 infection sites for each of the three replicates. Treatments: **CHA0**, plants obtained from seeds inoculated with CHA0 (10^6^ CFU/ml), **BABA**, plants soil-drenched with BABA (15 mM) 48h before rust infection, **Water**, plants mock-treated with sterile distilled water. Error bars indicate the standard error of the average values in 20 infection sites for each of three replicates. Asterisks indicate statistically significant differences in response to CHA0 or BABA treatments (Student’s t-test; α=0.05).

### Accumulation of H_2_O_2_ following the inoculation with leaf rust

The hydrogen peroxide released by plant tissue was measured between 0 and 72 hai with leaf rust in the control, the CHA0 and the BABA 15mM treatment. We monitored hydrogen peroxide with the DAB staining that produces dark-brownish dots (suppl. Fig. 2S). The surface of the dots was measured as the proportion of dark-brown pixels on the total leaf surface (Fig 5). Differences within the treatment became evident at 24 hai with higher H_2_O_2_ concentrations in the CHA0 and the BABA treated plants than in the control. Similarly, at 48 hai, in CHA0 and BABA treatments, the concentration of released H_2_O_2_ was significantly higher than in the control. At 72 hai, the accumulation of H_2_O_2_ in the CHA0 treatment dropped to the level of the control, while the BABA treatment increased to 12 % of the surface.

**Figure 5:**
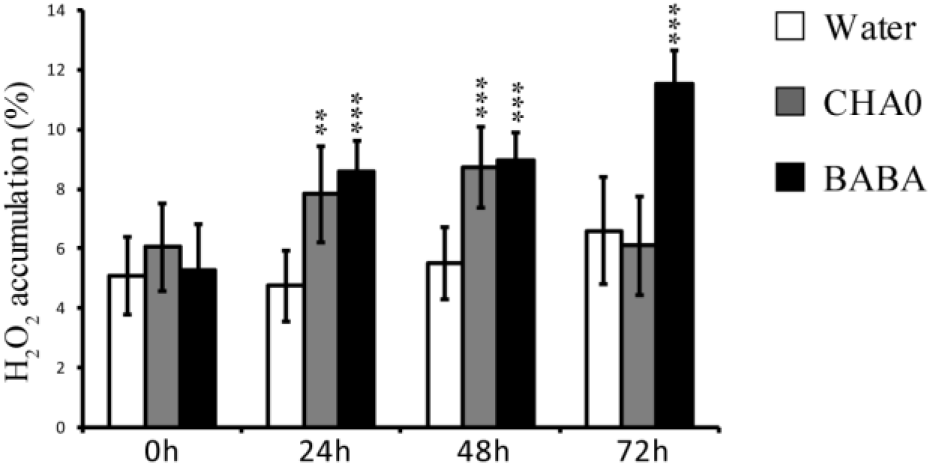
Hydrogen peroxide (H_2_O_2_) accumulation in wheat leaves in response to *P. triticina* infection. The generation of H_2_O_2_ was visible as dark-brown dots after DAB staining. The quantification based on the measurement of the dark-brown pixels on the photograph in proportion to the total leaf area. Treatments: **CHA0**, plants obtained from seeds inoculated with CHA0 (10^6^ CFU/ml), **BABA**, plants soil-drenched with BABA (15 mM) 48h before rust infection, **Water**, plants mock-treated with sterile distilled water. Error bars indicate the standard error for the average values of 10 replicates. Asterisks indicate statistically significant differences in response to CHA0 or BABA treatment (Student’s t-test; P =0.05).

## DISCUSSION

Induced resistance has proven to be a complementary control strategy potentially interesting for protecting wheat from foliar diseases (Görlach *et al.* 1996; Sharifi-Tehrani *et al.* 2009). Here, we confirmed the efficacy of both beneficial bacteria CHA0 and BABA.to induce resistance in wheat against leaf rust

We first assessed the physiological effect of both resistance inducers on wheat growth. Efficient root colonization by a given plant growth promoting bacterium is a prerequisite to exert a successful biocontrol effect on the host plant, either directly (*e.g.* disease suppression) or indirectly (*e.g.* ISR) (Lugtenberg and Kamilova 2009; Beneduzi *et al.* 2012). In our study, after seed inoculation, CHA0 was able to colonize wheat roots and more than 10^5^ CFU were recovered from 1g of root fresh weight. Our preliminary results showed that the initial concentrations used for seed inoculum (10^4^, 10^6^ and 10^8^ CFU/mL) did not affect final root colonization. The bacterial titer in wheat root was high enough for an effective plant protection as shown for soils suppressive to take-all of wheat and barley caused by *Gaeumannomyces graminis* var. *tritici* (Weller *et al.* 2007), Fusarium wilt of pea mediated by *Fusarium oxysporum* f. sp. *pisi* (Landa *et al.* 2002) and black root rot of tobacco (Stutz *et al.* 1986). Additionally, the growth promotion capacity of this stain was apparent with or without presence of leaf rust infections. In field experiments, a significant positive effect of beneficial soil organism application, including CHA0, on performance of wheat crop was observed especially when plants were under biotic stress (Imperiali *et al.* 2017). The observed plant growth promotion of CHA0 could be explained by production of phytohormones and the increase of nutrient availability to plants, in particular phosphate. CHA0 can solubilize mineral phosphate and improve plant growth in phosphate-limiting conditions (de Werra *et al.* 2009).

Recently, Thevenet *et al.* (2017) showed that BABA is a natural product in plants including wheat, but unfortunately, application of BABA can reduce plant growth in some plants (Cohen *et al.* 2016), At the relatively high concentration of 20 mM, BABA induced resistance but reduced the growth of the plant. The costs of induced resistance have also previously been linked to the reduction in plant growth (van Hulten *et al.* 2006; Heil 2007). Nevertheless, soil drenching with relatively low concentrations of BABA (15 mM) did not affect plant growth and reduced infection types in wheat seedlings infected with leaf rust, suggesting the possibility to optimize the BABA dose for an effective wheat protection against *P. triticina* with smallest impact on plant growth. Similarly, Luna *et al.* (2016) successfully identified feasible application methods of BABA by decreasing the concentration, which induced resistance in tomato against *Botrytis cinerea* without concurrent impacts on plant growth.

BABA is a well-recognized inducer of resistance against a broad spectrum of pathogens such as fungi, bacteria, virus and nematodes (Baccelli and Mauch-Mani 2016; Cohen *et al.* 2016). It is often applied as a soil drench (Hodge *et al.* 2005; Luna *et al.* 2016). Several studies demonstrate that BABA was effective when applied 1-3 days post-infection against a large spectrum of pathogens (Justyna and Ewa 2013). In the present, BABA was applied as a soil drench. 2 days before inoculation with leaf rust. The treatment significantly reduced leaf rust in wheat similar to results obtained with other rust species on wheat (Amzalek and Cohen 2007; Barilli *et al.* 2012). Inoculation with *P. protegens* strain CHA0 led to a specific reaction to infection with *P. triticina*: while mock-inoculated plants displayed many sporulating uredia (high infection type), plants with the bacterial treatment on the seeds had a mix of sporulating uredia and chlorotic and necrotic flecks, suggesting that *P. protegens* strain CHA0 partially reduced infection with *P. triticina* in the wheat seedlings.

Histopathological studies after induction of resistance were performed to identify the events that occur during pathogenesis, and ultimately lead to better understanding of the resistance mechanism.

The infection process of *P. triticina* in wheat plants has been well described (Bolton *et al.* 2008). During this infection process, the growth of the fungus can be interrupted at different phases. In principle, each of these phases can be affected by the action of resistance inducers. Our results regarding the infection events indicated that the pre-entry processes between plants treated with the two resistance inducers and control plants did not differ. In support of our results, reports show that the first steps of wheat rust infection (spore germination and appressoria formation) were not affected during the resistance implicated by the host plant (Wang *et al.* 2007; Orczyk *et al.* 2010). However, after penetration, distinct differences in fungal spread and host responses between CHA0 -and BABA-treated plants were observed. In BABA-treated plants, fungal penetration was strongly aborted 24 hai, based on the reduced percentage of substomatal vesicles. Moreover, the percentage of haustoria formed in BABA-treated plant was significantly reduced at 72 hai. This was also observed in durable resistance to leaf rust in the Brazilian wheat variety Toropi, where the number of haustoria formed was significantly reduced (Wesp-Guterres *et al.* 2013). Being effective at two important stages of fungal development could explain the infection type observed in BABA-treated plants, showing no to very small uredia formation surrounded by chlorotic flecks.

While, in CHA0-treated plants, small to medium uredia with no or low sporulation were apparent at the leaf surface and the number of pustules were reduced compared to infected control, this infection type could be explained by the fact that the CHA0 treatment was partially effective before haustorium formation and reduction in fungal penetration was observed at 24 and 48 hai. The successful penetrations generated a lower number of haustoria, giving later rise to small uredia.

To investigate the observed effect on fungal spread exerted by BABA and CHA0 treatment, assessment of callose deposition and hydrogen peroxide (H_2_O_2_) were performed. During fungal infection, callose can be deposited at infection sites, which provides a physical barrier preventing the penetration of a pathogen (Voigt 2016). Our results showed that callose depositions were mainly detected in guard cells. In support of our observations, Wang *et al.* (2015) demonstrated that the resistance response to *Puccinia graminis* f. sp. tritici is associated with callose deposition in the wheat guard cells. The increase of callose in BABA and CHA0-treated plants might restrict penetration and development of *P. triticina*, correlating with the increase of resistance in wheat seedlings against leaf rust. This defense mechanism of the plants is enhanced at the post-challenge primed state after perception of a stimulus from beneficial bacteria and BABA (Mauch-Mani *et al.* 2017). In addition to guard cells, callose was observed in the mesophyll cells of BABA-treated plants. This could also explain the high resistance observed compared to plants inoculated with bacteria. Same pattern was observed in defence mechanisms induced by *P. fluorescens* WCS417r and BABA against *Hyaloperonospora arabidopsis*. Both WCS417r and BABA prime for enhanced deposition of callose. However, more callose accumulated in BABA-treated plants (Van der Ent *et al.* 2009).

Reactive oxygen species (ROS) and especially H_2_O_2_ constitute a further important plant defence mechanism in interactions between plants and pathogens. We investigate H_2_O_2_ accumulation after infection with leaf rust in plants treated with BABA and CHA0. H_2_O_2_ accumulation was mostly detected in guard cells. At this site of penetration, an appressorium over the stomatal opening is generated. It seems likely that during the recognition or the formation of an appressorium, the generation of H_2_O_2_ in guard cells is induced possibly following secretion of rust elicitors. It might also be that mechanical forces during adhesion of the appressorium over the stoma elicit H_2_O_2_ generation in guard cells. In Arabidopsis, it was reported that H_2_O_2_ accumulation in guard cells is involved in the signal transduction during ABA-mediated stomatal closing (Sun *et al.* 2017). This could explain that accumulation of H_2_O_2_ in guard cells following recognition of leaf rust structures might be involved in stomatal closure, this being supported by the fact that we observed more accumulation of ABA in plants infected with leaf rust (data no shown), It had been reported that appressoria formation of *P. triticina* also caused stomata closure in wheat leaves (Bolton *et al.* 2008). Further studies shoe a correlation between H_2_O_2_ generation and hypersensitive reaction (HR) in resistance mechanism against wheat rust species (Wang *et al.* 2007; Orczyk *et al.* 2010; Serfling *et al.* 2016). In the current study, the accumulation of H_2_O_2_ due to both resistance inducers was observed 24 hai which corresponds to the beginning of haustorium generation. This suggests that H_2_O_2_ might initiate HR-defence mechanism. Our results are in line with the observation of Serfling *et al.* (2016) where HR was observed in mesophyll cells that were in contact with fungal haustorial mother cells at 24 hai, and the observed pre-haustorial resistance in the resistant accession PI272560 is due to an early HR of the first infected mesophyll cells. An HR accompanied by H_2_O_2_ accumulation also occurs in other interactions of plants with fungal parasites and causes non-host resistance to wheat stripe rust in broad bean (Cheng *et al.* 2012).

Plant-pathogen interaction can be modulated after induced resistance. Here we present a model for wheat-rust interaction (Fig. 6) where the infection of a host plant and growth of fungal structures have been interrupted at different phases in response to BABA or rhizobacteria-induced resistance. In control conditions, urediniospores of leaf rust were able to accomplish the infection cycle giving finally rise to an uredium with a normal size (Fig. 6A, left and right). In this case, callose accumulation in guard cells was not enough to prevent fungus penetration. Moreover, low generation of H_2_O_2_ was not able to initiate the required mechanisms to stop rust infection. While, in BABA and CHA0-treated plants fungal spread was differently affected (Fig. 6) with the exception of the pre-entry process where the spores germinated normally and appressoria were formed over the stomatal opening in both cases. In CHA0-treated plants, callose deposition in guard cells was highly elevated leading to an abortion of fungal penetration (Fig. 6B, left). However, when the fungus overcame the first barrier, callose deposition was not effective anymore. Here, we suggest that H_2_O_2_ accumulation can be accompanied by the activation of HR in some haustorium penetration sites which could partially stop fungal spread leading to the formation of small uredia (Fig. 6B, right), In the case of BABA, in addition to what we observed in CHA0 treatment, an accumulation of callose was noted in mesophyll cells. This could explain the high resistance observed after BABA treatment (Fig. 6C, left). Moreover, the high accumulation of H_2_O_2_ initiated HR in cells penetrated by rust haustoria and fungal spread totally stopped without any uredia formation (Fig. 6B, right).

**Figure 6:**
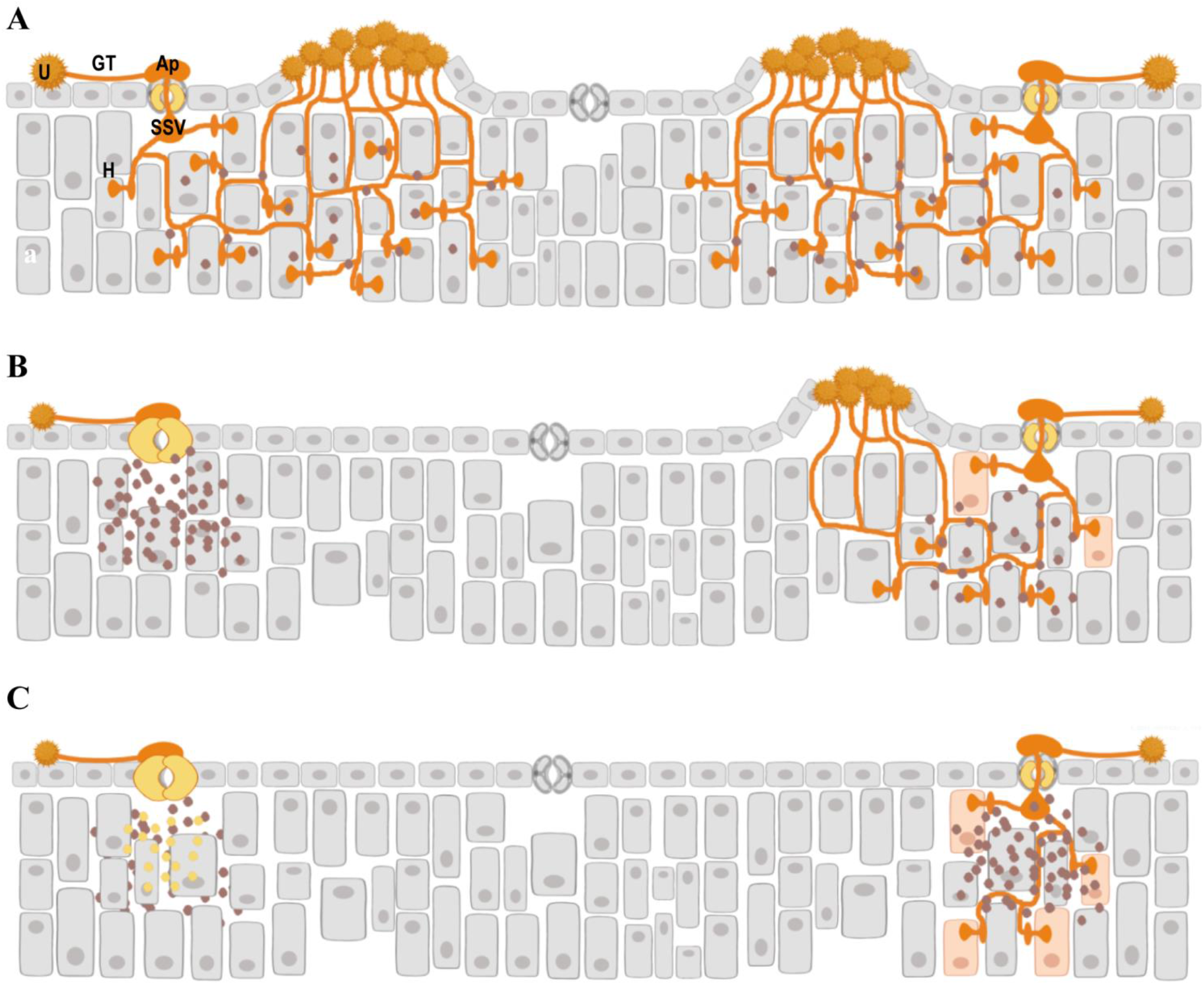
Diagram showing an overview of the fungal development and determined defense reactions of wheat to leaf rust infection under the effect of resistance inducers. **A**, compatible interaction between host and pathogen. In the un-treated plant, *P. triticina* overcomes the resistance mechanisms and is able to complete the infection cycle producing urediniospores (left and right). **B**, enhanced defense reactions in plants treated with CHA0; on the left, fungus penetration aborted after callose deposition in the guard cells, on the right, fungus spread partially but stopped after H_2_O_2_ accumulation and activation of HR in some haustorium penetration sites. Formation of small uredia without or with low spore production. **C**, enhanced defense reactions to leaf rust infection in BABA-treated plants; on the left, fungus penetration aborted after callose deposition in guard and mesophyll cells, on the right, fungus growth is totally blocked after accumulation of elevated quantities of H_2_O_2_ and HR activation in cells penetrated by rust haustoria. Fungal structures: **U**, urediniospore. **GT**, germ tube. **SSV**, substomatal vesicle. **Ap**, appressorium. **H**, Haustorium. Yellow dots are callose depositions. Brown spots present H_2_O_2_ generation.

The present study provides new insights into histological basis of BABA-and rhizobacteria-induced resistance against leaf rust of wheat showing the important role of callose deposition and H_2_O_2_ generation to prevent penetration and spread of leaf rust. Future studies will focus on expression analysis of some defense-related genes during the infection process of the fungus in order to underline differences and similarities in defence mechanisms induced by CHA0 and BABA.

## Acknowledgements

We thank Stefan Kellenberger, Agroscope Changins, Nyon, for technical support advice to handle the leaf rust pathogens. FB gratefully acknowledges the financial support by the Swiss Federal Commission for Scholarships for Foreign Students and BMM the financial support of the Swiss National Science Foundation, Grant No. 312 310030_160162.

## Supplementary material

**Table S1:** Simplified scheme of infection types of wheat leaf rust caused by *P. triticina* according to (Roelfs 1992)

| Response (class) | Infection type | Disease symptoms |
| --- | --- | --- |
| Immune | 0 | No uredia or other macroscopic sign of infection |
| Nearly immune | ; | No uredia but hypersensitive necrotic or chlorotic flecks present |
| Very resistant | 1 | Small uredia surrounded by necrosis |
| Moderately resistant | 2 | Small to medium uredia surrounded by chlorosis or necrosis |
| Moderately susceptible | 3 | Medium-sized uredia that may be associated with chlorosis |

**Figure 1S:**
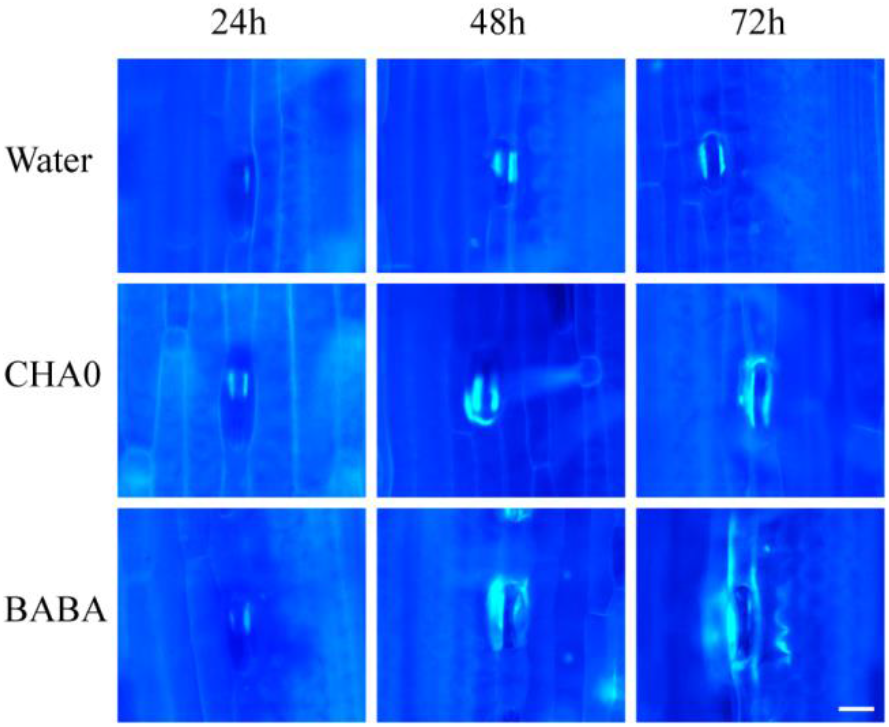
Localization of callose at 24, 48 and 72 hai by *P. triticina* in wheat leaves treated with CHA0 or BABA. Photographs show stained leaves (Aniline-blue) exposed to UV light. Treatments: **CHA0**, plants obtained from seeds inoculated with CHA0 (10^6^ CFU/ml), **BABA**, plants soil-drenched with BABA (15 mM) 48h before rust infection, **Water**, plants mock-treated with sterile distilled water. Bar 20 μm.

**Figure 2S:**
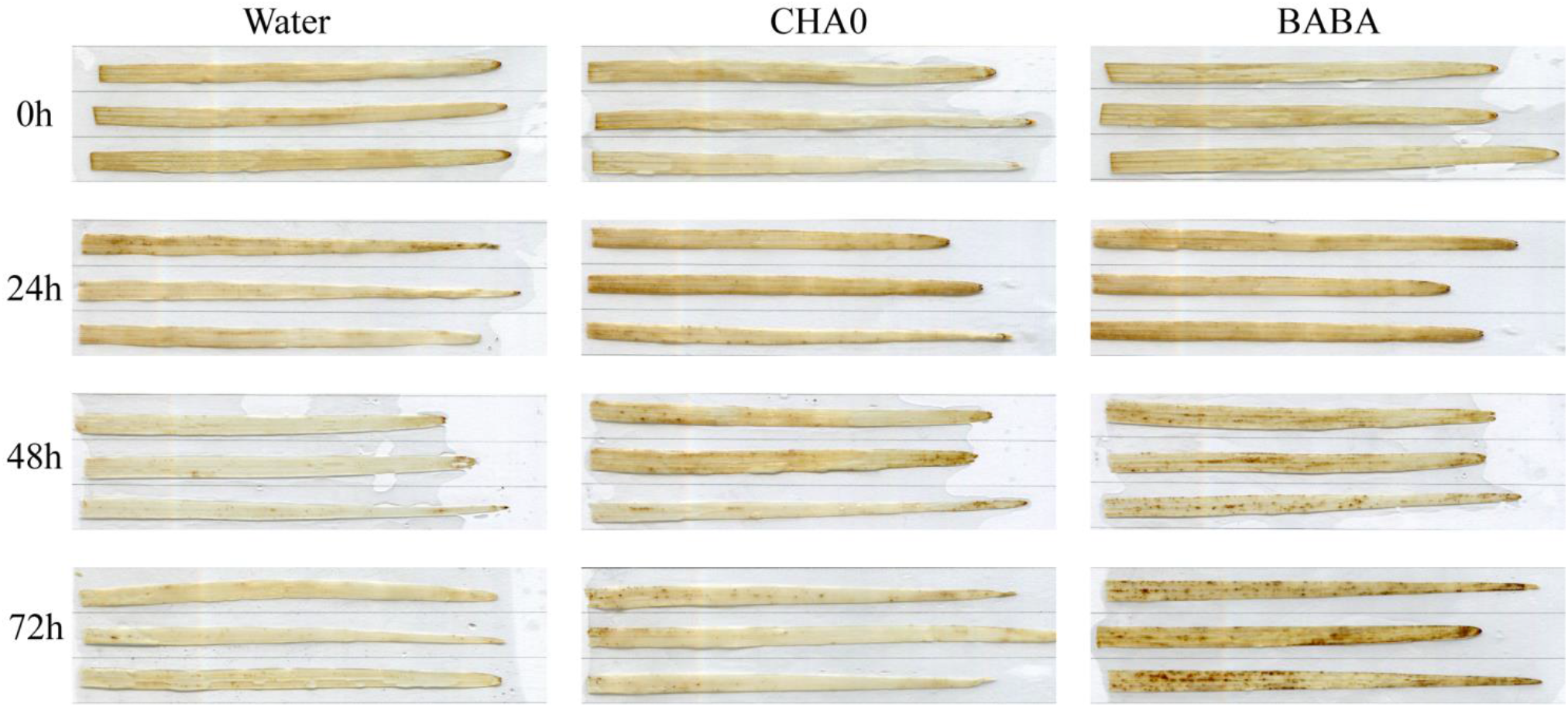
*In situ* detection of hydrogen peroxide (H_2_O_2_) using DAB staining at 0, 24, 48 and 72 hai by *P. triticina* in wheat leaves treated with CHA0 or BABA. Images were obtained by scanning at 1.200 dpi the stained second leaf. Treatments: **CHA0**, plants obtained from seeds inoculated with CHA0 (10^6^ CFU/ml), **BABA**, plants soil-drenched with BABA (15 mM) 48h before rust infection, **Water**, plants mock-treated with sterile distilled water.

